# Correlated seed failure as an environmental veto to synchronize reproduction of masting plants

**DOI:** 10.1101/171579

**Authors:** Michał Bogdziewicz, Michael A. Steele, Shealyn Marino, Elizabeth E. Crone

**Affiliations:** Department of Systematic Zoology, Faculty of Biology, Adam Mickiewicz University, Umultowska 89, 61-614 Poznań, Poland; CREAF, Cerdanyola del Valles, 08193 Catalonia, Spain; Department of Biology, Wilkes University, Wilkes-Barre, PA 18766, USA; Department of Biology, Tufts University, 163 Packard Ave, Medford, MA 02155, USA

**Keywords:** density-dependent pollination success, environmental veto, mast seeding, Moran effect, resource budget model, seed production, *Quercus alba*, *Quercus rubra*

## Abstract

Highly variable, synchronized seed production, called masting, is a widespread reproductive strategy in plants. Resource dynamics, pollination success, and, as described here, environmental veto, are possible proximate mechanisms of masting. We extended the resource budget model of masting with correlated and uncorrelated reproductive failure, and ran this model across its parameters space. Next, we parametrized the model based on a 16-year seed production data for red (*Quercus rubra*) and white (*Q. alba*) oaks. Simulations showed that resource dynamics and reproduction failure produce masting even in the absence of pollen coupling. In concordance, in both species, among-year variation in resource gain and correlated reproductive failure were necessary and sufficient to produce masting. Environmental variation is a form of reproduction failure caused by environmental veto that may drive large-scale synchronization without density-dependent pollen limitation. Reproductive-inhibiting weather events are prevalent in ecosystems, suggesting that these described mechanisms likely operate in many masting systems.

## Introduction

Synchronized, high inter-annual variability in seed production, called mast seeding, characterizes reproduction of numerous plant species (Kelly, 1994). Our understanding of the proximate mechanisms synchronizing reproduction among individuals and forcing plants to produce seeds irregularly is incomplete (Pearse *et al.*, 2014; Pérez-Ramos *et al.*, 2015, Koenig *et al.*, 2016; Pearse *et al.*, 2016). Weather could synchronize reproduction among plants by providing an environmental cue for mast flowering (cf. Kelly, 1994; Kelly *et al.*, 2013). Pollen limitation has also been implicated in synchrony. First, successful pollination may depend on weather conditions, creating a Moran effect, i. e. weather conditions affecting effectiveness of pollen transfer among plants (Crone & Rapp, 2014; Pearse *et al.*, 2016). Second, density-dependent pollen limitation can create feedbacks that synchronize plants within populations (Satake & Iwasa, 2000). The notion that pollination is more efficient in high-flowering years has been around for some time (e.g., Stearns, 1976; Nilsson & Wastljung, 1987; Norton & Kelly, 1988; Smith *et al.*, 1990, Kelly *et al.*, 2001). However, its importance as a synchronizing mechanism, as opposed to a fitness benefit alone, has only recently been appreciated (Satake & Iwasa, 2000; Satake & Iwasa, 2002; Crone *et al.*, 2005). Although researchers agree that seed production is resource limited, there is no consensus about the exact mechanisms that produce masting (Crone & Rapp, 2014; Pearse *et al.*, 2016), and there are few studies that simultaneously test different drivers of masting.

The resource budget (RB) model formalized the hypothesis that mast-seeding results from resource storage and allocation to reproduction, in combination with synchronizing mechanisms (Isagi *et al.*, 1997; Satake & Iwasa, 2000). The RB model assumes that plants need to accumulate a threshold level of resources to initiate reproduction. If this threshold cannot be reached in one season, then plants accumulate resources over several years, which translates into intermittent seed production by individual plants (Isagi *et al.*, 1997). Although resource dynamics cause intermittent, supra-annual, seed production at the individual level, each plant could maintain its own schedule of high and low years of seed production, in the absence of other synchronizing factors (Isagi *et al.*, 1997; Satake & Iwasa, 2000). The hypothesis of synchrony via density-dependent pollen coupling has been well developed in theory (Satake & Iwasa, 2000; Satake & Iwasa, 2002a,b; Venner *et al.*, 2016), and demonstrated in a handful of case studies (Crone *et al.*, 2009, Rapp *et al.*, 2013, Abe *et al.*, 2016). Synchrony can also be achieved by weather cues (Rees *et al.*, 2002; Monks *et al.*, 2016). Specifically, although simple among-year variation in resource gain is not sufficient to synchronize seed production (Satake & Iwasa, 2002a,b), weather cueing may be effective at synchronizing plants when it functions as an environmental veto, in which certain weather events prevent nearly completely prevent flower or seed set over a significant geographic area (Crone & Rapp, 2014; Pearse *et al.*, 2016, Abe *et al.*, 2016).

In this study, we explore the environmental veto hypothesis in greater detail, using a combination of mathematical models and patterns of acorn production in two species of oaks (*Quercus* spp.) of eastern North America. Generally, acorn production in oaks is thought to be resource-limited (Sork *et al.*, 1993, Koenig *et al.*, 2003, Pesendorfer *et al.*, 2016). However, the drivers of synchrony are less well understood, and it is not clear whether synchrony is due to environmental variability or pollen coupling (Liebhold *et al.*, 2004, Koenig & Knops 2014, Koenig *et al.*, 2016, Bogdziewicz et al. 2017). Lack of a direct relationship between flower abundance and acorn crop has been reported in many (Sharp & Sprague, 1967; Sork *et al.*, 1993; Pearse et al. 2015), but not all (Sork *et al.*, 1993; Sork & Bramble, 1993) studies of oaks. However, in numerous oak species, spring temperature correlates positively with acorn production (Sork & Bramble, 1993; Koenig & Knops, 2014), and warm springs enhance synchrony of flowering, which correlates with crop size (Koenig *et al.*, 2015, Bogdziewicz *et al.*, 2017). In contrast, low temperatures desynchronize flowering and lead to pollination failure (Koenig *et al.*, 2015, Bogdziewicz *et al.*, 2017). Another environmental forcing mechanism that can veto reproduction and enhance synchrony in oaks is catastrophic weather events, e.g. late spring frosts that kill flowers (Sharp & Sprague, 1967; Cecich & Sullivan, 1999).

Our study involves three approaches. First, we explore the role of reproductive failure (e.g. driven by environmental veto) as a driver of masting using an extension of the RB model. Specifically, we simulate reproductive failure that is either perfectly correlated among individuals within a population (cf. Abe *et al.*, 2016), or affects individuals independently, affecting overall variation of seed production at the individual and population level. By doing so, we explore how extended RB model parameters interact in creating plant masting behavior. Second, we analyze three 16-year long time series of seed production by individual trees of two wind-pollinated, self-incompatible oak species: northern red (*Quercus rubra*) and white (*Q. alba*) oaks, and characterize temporal patterns of seed production in these six populations (three of each species). Finally, we extend the general approach of Rees *et al.*, (2002) and Crone *et al.*, (2005), and use these time series to parameterize RB models with various combinations of synchronizing factors (density-dependent pollen coupling, and environmental veto due to correlated seed failure). We use these empirically-based models to simulate dynamics of seed production through time, and compare seed production in simulated time series to patterns observed in the field populations.

## Methods

### 1. Resource budget model with seed failure

Our analysis is based on the RB model derived from Isagi *et al.*, (1997) and Satake & Iwasa (2000), and slightly modified by Rees *et al.*, (2002) and Crone *et al.*, (2005). The model assumes that flowering is conditional on whether a resource threshold is reached by an individual plant (*D*_*i*,*t*_ ≤ *C* for not flowering, *D*_*i*,*t*_ ≥ *C* for flowering plants), where *C* is the minimum resource threshold needed for reproduction, *D_t_* is the amount of the internal resources of plant *i* in year *t*, and each plant *i* gains resources *P*_*i*,*t*_ in year *t*. Once the level of resources exceeds the threshold, a plant initiates reproduction and allocates an amount of resources proportional to the excess into flowering, and depletes resources by an amount *A*(*D*_*i*,*t*_ – *C*), where *A* is the resource cost of reproduction (fruiting-to-flowering cost ratio, the depletion coefficient). In the presence of density-dependent pollen coupling, seed production and resource depletion depend on the number of flowering plants; specifically, resource depletion becomes
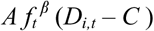
where *f_t_* is the proportion of flowering plants in population in year *t* (as a surrogate of pollen availability), and *β* scales the strength of pollen coupling. Small *β* indicates high pollination efficiency (low pollen coupling), while high *β* indicates that pollination efficiency in an individual plant strongly depends on the flowering of the others in the population (Satake & Iwasa, 2000). These assumptions lead to the following model of stored resource dynamics through time, where all parameters are as defined above:

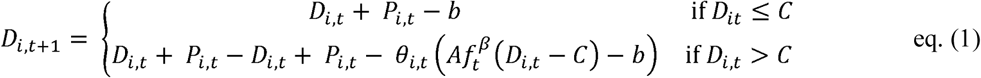

We extended the RB model to include an environmental veto that prevents reproduction even though resources are available. Specifically, *θ*_*i*,*t*_ is the probability that a plant in year *t* will experience unfavorable environmental conditions that prevent seed production (Abe *et al.*, 2016), and *E*_*i*,*t*_ ≥ *θ*_*i*,*t*_ is a random number uniformly distributed between 0 and 1:

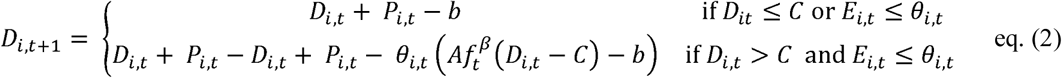

We explored the properties of this model by running two sets of simulations for a population of 200 plants over 1000 steps (years). In the first set, the probability of reproduction failure for a given year was identical for all plants within the population (same *E*_*i*,*t*_ for all plants in year *t*, with a constant value of *θ* for all years. We ran simulations with *θ* from 0-0.25 (26 models with probability increasing from 0 to 0.25 by 0.01 increments). In the second set, we explored the same range of probabilities but we let plants to fail or not independently from each other (*θ*_*i*,*t*_ = *θ* for all plants, but with a different *E*_*i*,*t*_ drawn for each plant, i.e., a binomial distribution of failure with the same probability of failure for all plants within a year; hereafter uncorrelated probability of failure). Again, we ran the uncorrelated failure model with *θ* from 0-0.25 (26 scenarios with probability increasing from 0 to 0.25 by 0.01 increments). We ran both sets of models across a range of parameters space for the depletion coefficient (*C* in eq. 1, from 0.5 to 3.5, which represents low and high proportional costs of fruits to flowers production, respectively) and three levels of pollen coupling *β* (0, 0.5, and 1).

We used the last 500 years of these simulations to evaluate patterns of seed production. With these data, we characterized simulation results using four metrics of variation and synchrony in seed production, commonly used in studies of mast seeding (Herrera *et al.*, 1998; Koenig *et al.*, 2003; Crone *et al.*, 2011): variability in seed production of individuals (CV_i_), population-level variability of seed production (CV_p_), lag-1 autocorrelation of seed production at the population level (ACF1), which is a measure of the tendency of populations to alternate between years of high and low seed production, and synchrony of seed production by plants (*S*), as measured by the average pairwise correlation of seed production of individual plants in a population through time.

### 2. Field studies of acorn production in oaks

#### Study system

Both red and white oaks are dioecious, i.e., individual plants produce separate male and female flowers. Red and white oak belong to two different sections of oaks (*Lobatae* and *Quercus*, respectively). Because of this difference, the two species differ in length of time between flower production and acorn maturation (Sork *et al.*, 1993). White oaks produce flowers in spring, which are fertilized and develop into mature fruit in the same year as they were pollinated. Red oak produce flowers that are pollinated during the spring of one year, but the fruit mature the following year. Both species develop flower primordia in the year prior to flowering (Sork *et al.*, 1993), which means that the decision to produce flowers is made one year before the plant begins to allocate resources to reproduction (1.5 years before seed maturation) in white oaks, and approximately two years before resource allocation (2.5 years before seed maturation) in red oaks. However, for both species, acorn filling occurs in the year when acorns are shed. Therefore, it is plausible for both species that seed production depends on resources at the start of the growing season in which acorns are produced, as assumed by resource budget models.

#### Field monitoring

We monitored acorn production over 16 years at three study areas in eastern Pennsylvania, USA: two in Luzerne County (Steele, 41.135966, -75.992672; and Venesky 41.160088, -75.932110) and a third in Skhuylkill County (Hawk Mountain, 40.636821, -75.981969) (see detailed description in Moore *et al.*, 2007). We monitored acorn production by individual oak trees by placing two seed traps under each of at least 15 individuals of each species at each site from 2001-2016.

#### Data analysis

We characterized time series for each species at each site using the four metrics described above (see *Methods 1. Resource budget model with seed failure*): population- and individual-level coefficients of variation (CV_p_, and CVi), lag-1 autocorrelation (ACF1), and synchrony (S). We used these metrics to compare observed time series of seed production to simulation model outputs.

### 3. Empirical resource budget models

#### Parameter estimation

To reconstruct a plant’s stored resources from acorn production data we followed the general approach developed by Rees *et al.*, (2002). In brief, this analysis involves first, fitting a linear model of cumulative reproduction (summed acorn count) versus cumulative years (as surrogate of resource acquisition over time) for each oak species. We used generalized linear mixed models (GLMMs) implemented via ‘lme4’ package (Bates *et al.*, 2015) with tree ID as random intercept and year as random slope. The random effect structure allowed fitting a unique intercept for each plant which estimates stored resources of a plant at the beginning of the study (Rees *et al.*, 2002), while random slope allowed heterogeneous resource acquisition of individuals over time (Crone *et al.*, 2005). Residuals of these two models (one for each species) are an estimate of stored resources of each individual plant has in each year (Rees *et al.*, 2002; Crone *et al.*, 2005).

To estimate pollen coupling and reproduction failure, we analyzed acorn production by each individual tree in each year, as a function of its estimated stored resources (Rees *et al.*, 2002) and the proportion of reproductive plants in each population (Crone *et al.*, 2005). To incorporate the possibility of seed failure, we used zero-inflated negative binomial (ZINB) mixture models implemented via ‘*pscl*’ package (Zeileis *et al.*, 2008; Jackman, 2015). Mixture models analyze the data in two parts with the first one being a binomial process that estimates the probability of measuring a zero (here, no acorns produced by a tree), while in the second process acorn count is modeled by negative binomial model; in other words, the analysis separates “true” failure from zero counts due to sampling variation (Zuur *et al.*, 2009). Within this framework, we constructed statistical models to match the exact structure of the theoretical model (eq. 2). Specifically, we rearranged the modified RB model to predict seed production dynamics, *S*_*i*,*t*+1_, as a function of stored resources, *D*_*i*,*t*_:

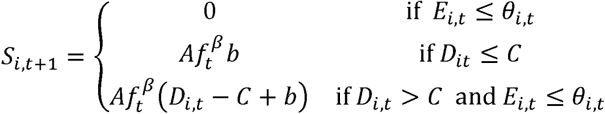

where *b* is a coefficient allowing for a small amount of flowering in years when plants are below threshold (cf. Rees *et al.*, 2002), and all other parameters are as described in equations 1 & 2. To demonstrate the relationship fits to a 0-inflated binomial model, we transform the dynamics to a log scale (consistent with use of a log link in a ZINB model):

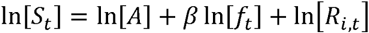

where

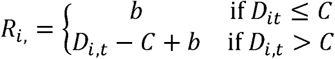

From these relationships, it is clear that the intercept term for the count part of a ZINB model is an estimate of In[*A*], the slope with respect to In[*f_t_*] is an estimate of *β*, and the model should include an offset of ln[*R*_*i*,*t*_].

We explored model structures representing different hypotheses about environmental correlation and pollen coupling. Environmental correlation of seed failure was included by making the zero term in the model a function of site and/or year. Pollen coupling was included by including ln[f_t_] in the model (or not). Specifically, we first used the full model structure to determine an appropriate model for environmental correlation of seed failure: year effects, site effects, or site + year effects (site × year models did not converge). These were implemented by changing the predictor variables for the 0-inflation term. Next, we evaluated four models representing all possible combinations of correlated seed failure (or not) and pollen coupling (or not). To implement these models in the *pscl* package, we iterated over possible values of *b* and *C* to find maximum likelihood values for each model structure.

Because red oak flowers develop one year before fertilization, we also fit models to this species using a modified RB model in which seed production in one year is determined by resource status at the end of the previous year (see Online Appendix for details).

#### Simulations of seed production

We used parameters estimated for each of the four model structures (RB alone, RB + pollen coupling, RB + correlated seed failure, RB + pollen coupling + correlated seed failure) to simulate time series of acorn production for populations of each species. We simulated seed production by populations of 20 trees for 500 years. We used the last 15 steps (years) of each simulation to calculate masting parameters (CV_p_, CV_i_, S, and ACF1, as described above, see *Methods: General theoretical models*).

For each model structure and species, we conducted two sets of simulations. The first included seed failure as the only source of environmental variation. Specifically, we calculated a logit-scale mean and standard deviation in the probability of seed failure using the annual estimates of the probability of 0’s in the ZINB model. For both species, there were significant effects of site as well as year on the probability of 0’s (see *Results*), so we used the average mean and standard deviation across sites for simulations. Then, in each year, we first sampled the probability of seed failure for that year, then used a binomial distribution to determine whether each individual tree experienced seed failure. For models with uncorrelated seed failure, we used the same probability for all years, but still included binomial variation in failure among individuals.

We also conducted a second set of simulations in which we included additional environmental stochasticity in costs of reproduction, based on the variance of the count term (seeds per stored resource) of the ZINB model. Variation in costs of reproduction and annual resource gain are mathematically confounded in the RB model (Satake & Iwasa, 2002a), so this term combines both sources of variation. For this model, we separated the negative binomial distribution for the count term into its component Poisson and gamma distributions. Using the negative binomial distribution as parameterized in the zeroinfl function, the overdispersion parmeter, *θ*, is the reciprocal of the variance of the component gamma distribution, so the appropriate gamma distribution has shape parameter *θ* and scale parmeter 1/*θ*, based on the value of *θ* for each model structure. We sampled from this gamma distribution to obtain average seed set per resource in each year, then determined seed set of each tree by sampling from a Poisson distribution with the annual mean.

## Results

### 1. Resource budget model with seed failure

Our simulations showed that resource dynamics together with correlated reproduction failure can produce masting behavior in plants even in the absence of pollen coupling. The CV_p_ of seed production increased with the probability of correlated failure for all values of resource depletion coefficient and exceeded CV_p_ = 1for failure probabilities above 10% (Fig. 1). The highest CV_p_ was produced in populations with both high pollen coupling and high probability of correlated reproduction failure (CV_p_ ranged 2-3 for depletion coefficient > 2 and reproduction failure > 15%, Fig. 1). In contrast, uncorrelated seed production failure broke down masting and led to small CV_p_ for all values of depletion coefficient and *β* (most simulations resulted in CV_p_ < 0.5, Fig. 1).

**Fig. 1.**
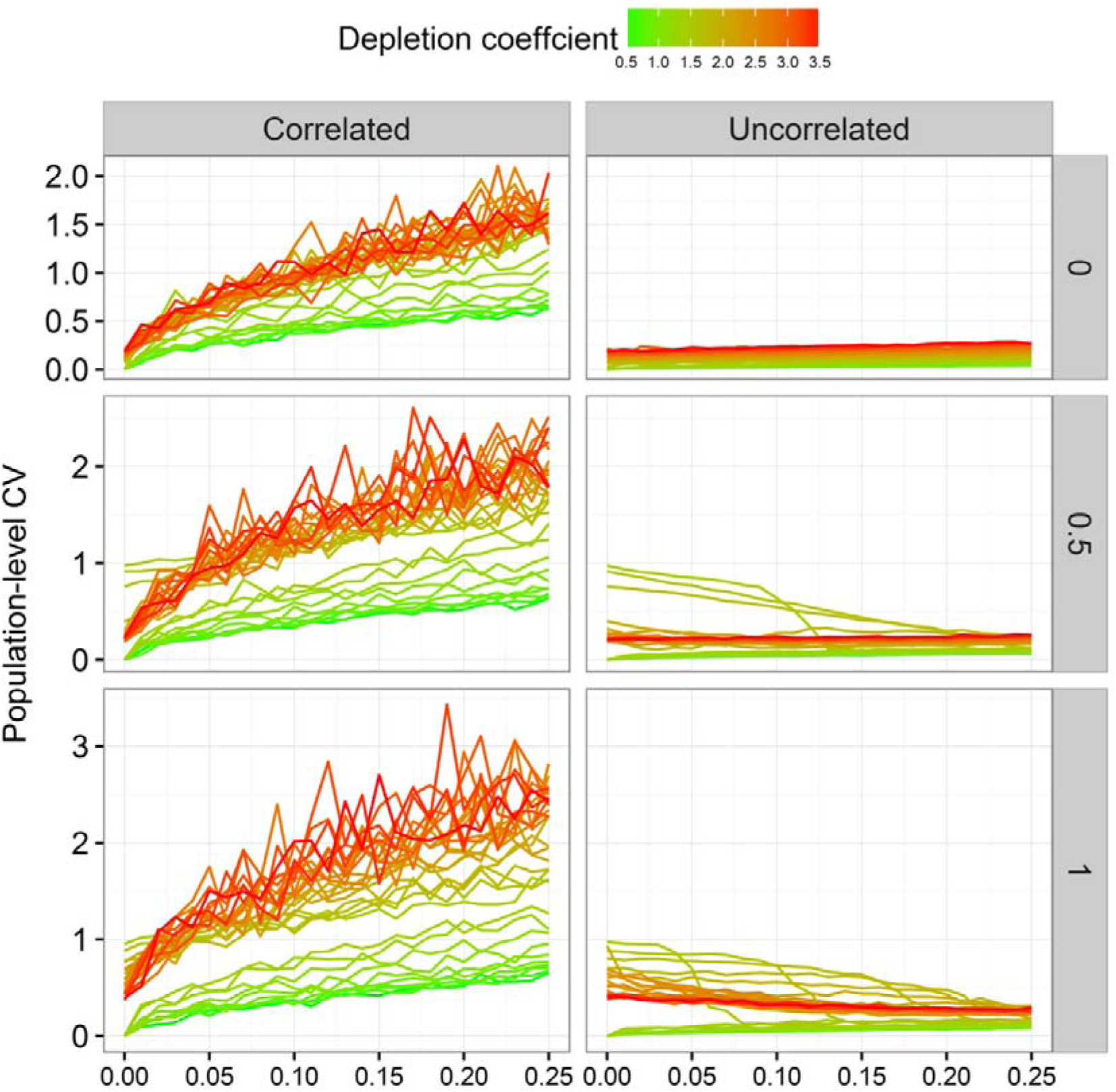
Simulated seed production dynamics (CV_p_) across different values of probability of correlated and uncorrelated seed production failure, and strength of pollen coupling, *β*: 0 (low), 0.5 (moderate), and 1 (high). Seed production is resource-dependent with depletion coefficient ranging from 0.5 – 3.5.

These contrasting effects of correlated and uncorrelated failure probability on CV_p_ were caused by different effects of these on seed production synchrony (S). Correlated reproduction failure increased synchrony, especially for low values of depletion coefficient (Fig.1S). The strength of the positive relationship between correlated reproduction failure and *S* decreased with increasing values of depletion coefficient (Fig. 3S). This effect was attenuated by pollen coupling, and the synchrony was highest (S > 0.5) for *β* = 1 and correlated failure probability above 15%. In contrast, uncorrelated seed production failure desynchronized reproduction within population for all values of depletion coefficient and *β*, and scaled down S to 0 for most simulations (Fig. 3S).

In case of the other masting metrics, ACF1 decreased with increasing probability of seed failure (both un- and correlated) for intermediate values of depletion coefficient, and had no effect for higher depletion coefficient (Fig 2S). Moreover, in the presence of pollen coupling correlated (but not uncorrelated) reproduction failure increased CV_i_ (Fig. 3S). In summary, the correlated reproduction failure enhances mast seeding behavior (CV_p_) through increasing CV_i_ and synchrony (S) of seed production.

### Field studies of acorn production in oaks

Time series of both species were typical of mast seeding trees. All six populations had CV_p_ > 1; for white oak, the CV_p_’s were 1.53, 1.82 and 1.64, and for red oak the CV_p_’s were 1.18, 1.66 and 1.67. Individual-level CV’s were higher than population-level CV’s: CVi = 2.02, 2.63, and 2.19 for white oak and CVi = 1.71, 1.81, and 2.74 for red oak. Synchrony (the correlation among individual trees in acorn production through time) of individuals within populations was consistently high and positive; S = 0.58, 0.20 and 0.36 for white oak, and S = 0.45, 0.71, and 0.41 for red oak. Synchrony of individuals among populations was weaker then synchrony within populations, but still positive (white oaks S = 0.33, red oaks S = 0.27). Synchrony among individuals of the different species within each site was consistently lower than synchrony among individuals of the same species (S = 0.31, 0.33 and 0.35 for red oak-white oak synchrony at each site). Time series also consistently had negative lag-1 autocorrelations; -0.10, -0.23, and -0.11 for white oak and -0.01, -0.13, and -0.14 for red oak.

However, in one case AC1 = -0.01 (for red oak at Hawk site), this metric is similar to the expectation for short, uncorrelated time series (E. Crone, unpubl. analyses).

### Empirically-based models: parameter estimation

GLMMs of cumulative acorn production through time showed higher acorn production in red oaks than white oaks (slope coefficients of 3.19 ± 0.39 (SE) and 2.40 ± 0.32 (SE), respectively), as well as substantial variation among individual trees for both species (SD of random effect of slope coefficient × individual tree = 2.86 ± 0.26 (SE) and 2.32 ± 0.18 (SE) respectively). Residuals from these models show the expected pattern of higher seed production in years following high residuals, with a noticeable threshold for reproduction and some evidence of acorn failure (zero’s in years when plants had sufficient resources to reproduce; Fig 2).

**Figure 2.**
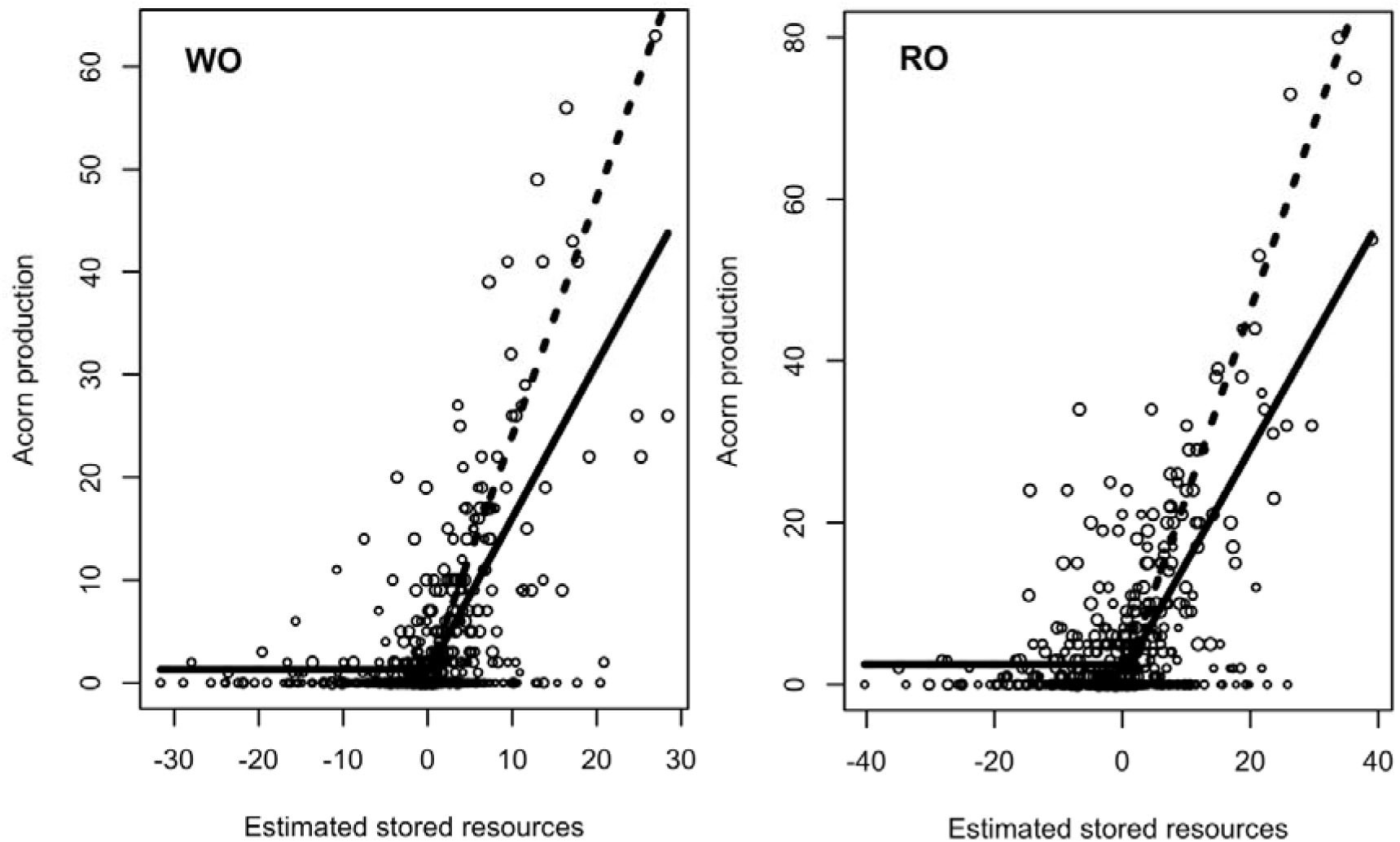
Acorn production in white (WO) and red (RO) oaks as a function of estimated stored resources. Dashed line indicate estimates obtained from ZINB model with pollen coupling and correlated seed production failure, and solid line indicates model with correlated reproduction failure only. Diameter of circles indicates the number of reproducing trees at each site in each year. Models with higher slope fit the data better (see Results), which indicates higher per resource acorn production with increasing number of reproducing individuals within population, i.e. pollen coupling.

For white oaks, seed failure (i.e., the probability of 0’s in the ZINB models) differed significantly among sites (χ^2^ = 20.8, d.f. = 2, *P* = < 0.001), with the highest probability of seed failure at Steele (0.81), followed by Venesky (0.58), and Hawk (0.53). Failure also differed significantly among years (χ^2^ = 74.7, d.f. = 14, *P* < 0.001). Similarly, for red oaks, seed failure differed significantly among sites (χ^2^ = 28.5, d.f. = 2, *P* = <0.001), with the highest probability of seed failure at Venesky (0.87), followed by Steele (0.72) and Hawk Mountain (0.60). Failure also differed significantly among years (χ^2^ = 55.0, d.f. = 14, *P* = <0.001). Therefore, for both oak species, we use the model with both main effects for all further analyses.

For both species, the best models of acorn production included both correlated seed failure and density-dependent pollen coupling (ΔAIC ≥ 34.3 and ΔAIC ≥ 30.7 for white and red oaks, respectively; Table 1). In this model structure, the pollen coupling coefficient, *β*, was higher for red than white oak, indicating stronger pollen coupling in the former (Table 1). Similarly, for red oak the second-best model structure (closest AIC) was for the model with pollen coupling only, whereas the second-best model for white oaks was the model with correlated seed failure only (Table 1). The proportion of years with seed failure was higher and more variable among years for white oaks than for red oaks (Table 1).

**Table 1.** Model selection statistics and parameter estimation based on zero-inflated negative binomial models of acorn production in white and red oaks vs. estimated stored resources. RB indicates resource-only models, RB+PC is resource-based and pollen coupling model, RB+EC is resource-based and environmental correlation model, and RB+PC+EC is resource-based, pollen coupling and environmental correlation model.

|  | White oaks |  |  |  | Red oaks |  |  |  |
| --- | --- | --- | --- | --- | --- | --- | --- | --- |
| Model | RB | RB+PC | RB+EC | RB+PC+EC | RB | RB+PC | RB+EC | RB+PC+EC |
| Model AIC | 2081.7 | 1896.7 | 1879.2 | 1844.8 | 2471.0 | 2265.9 | 2332.2 | 2234.0 |
| Parameter estimates: |  |  |  |  |  |  |  |  |
| Depletion coefficient<sup>a</sup>, A | 1.57 | 3.25 | 1.51 | 2.32 | 1.36 | 2.99 | 1.39 | 2.30 |
| Pollen coupling<sup>b</sup>, b | NA | 1.48 | NA | 0.75 | NA | 1.57 | NA | 1.14 |
| Mean proportion of years with no reproduction<sup>c,e</sup> | 0.507 | 0.11 | 0.53 | 0.51 | 0.51 | 0.17 | 0.32 | 0.31 |
| SD proportion of years with no reproduction<sup>d,e</sup> | NA | NA | 0.32 | 0.29 | NA | NA | 0.30 | 0.26 |
| Logit-scale mean<sup>c</sup> | 0.03 | -2.12 | 1.72 | 1.79 | 0.05 | -1.57 | -1.87 | -1.20 |
| Logit-scale SD<sup>d</sup> | NA | NA | 6.18 | 6.05 | NA | NA | 4.16 | 1.73 |
| Resource threshold for reproduction<sup>f</sup>, C | 0.4 | 0.4 | 0.2 | 0.4 | 1.0 | 0.6 | 1.0 | 0.6 |
| Reproduction below threshold<sup>g</sup>, b | 0.9 | 1.1 | 1.3 | 1.6 | 2.5 | 2.0 | 2.5 | 2.7 |
| Among-year variation in seed production, 1/$\theta$ | 1.555 | 2.037 | 0.962 | 0.774 | 1.115 | 1.588 | 1.104 | 0.924 |

<sup>a</sup>Back-transformed ( $\exp()$ ) intercept for count term in ZINB model
<sup>b</sup>slope of $\ln(\text{seed production})$ vs. $\ln(\text{proportion plants flowering})$ for count term in ZINB model
<sup>c</sup>Average proportion of 0 years in ZINB model; averaged across sites where site is statistically significant
<sup>d</sup>standard deviation of year-specific coefficients for the proportion of 0 years in ZINB model; averaged across sites where site is statistically significant
<sup>e</sup>Calculated by, first, back-transforming values, and then calculating the mean and SD of the back-transformed values
<sup>f</sup>Break-point in piecewise regression for count term in ZINB model
<sup>g</sup>Reproduction below the threshold for count term in ZINB model (estimated as part of an offset term)

For red oaks, parameter estimates were similar for variants of the “standard” RB model with seed production based on resources at the beginning of that year, and for the lagged RB model, with seed production based on resources at the beginning of the previous year (Table S1). Therefore, in the rest of this paper, we explore only the unlagged models.

### Empirically-based models: stochastic simulations

As expected from general theory, the RB model with no synchronizing mechanism predicted highly variable seed production by individual trees (CV_i_ = 2.07 and 1.65 for white and red oaks, respectively, Figure 3), but no synchrony at the population level (S = 0 for both species, and CV_p_ = 0.45 and 0.36 for white and red oaks, respectively). Models with pollen coupling but no source of environmental forcing had more synchronous reproduction (S = 0.28 and 0.22 for white and red oaks, respectively), but were not more variable at the population level (CV_p_ = 0.16 for white and 0.22 for red oaks).

**Figure 3.**
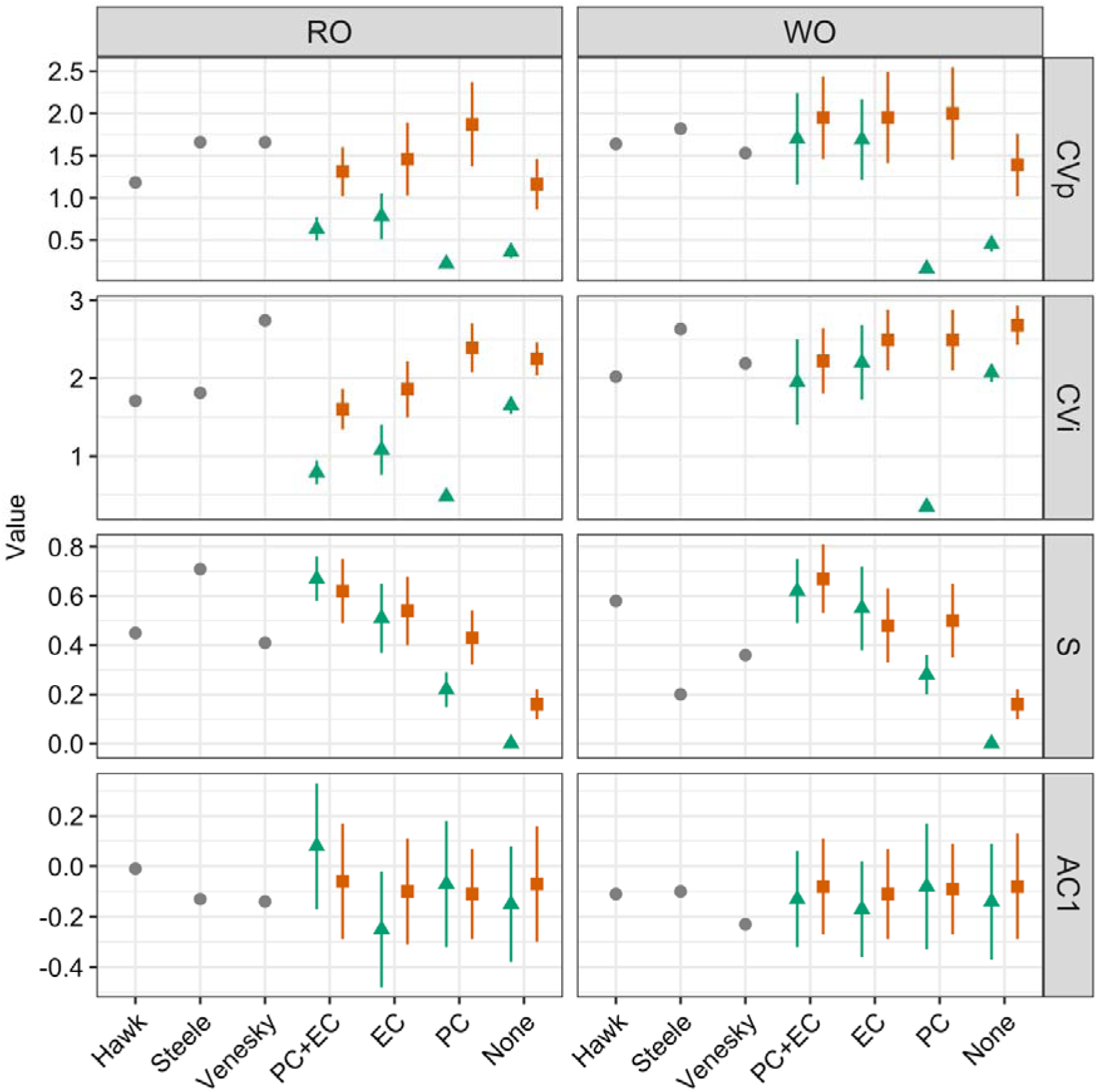
Masting metrics for the red (RO) and white (WO) oak populations (sites Hawk, Steele, Venesky; grey circles), and for simulated seed production dynamics. Green triangles indicate seeding dynamics from deterministic resource budget models (i.e. resource acquisition constant over years), while orange squares from stochastic models (i.e. variable resource gain over years). PC + EC indicate resource-based models with pollen coupling and correlated environmental failure, EC with correlated environmental failure, and PC with pollen coupling as synchronizing mechanisms, while ‘None’ is for models with either PC or EC. Simulations are based on the extended resource budget model (cf. eq. 2), and where parametrized using the estimations from zero-inflated negative binomial models of acorn production in both species (cf. Table 1).

For red oaks, adding correlated seed failure to simulations increased the CV_p_, but not enough to reproduce dynamics (CV_p_ = 0.63 and 0.78 for simulated time series from models with correlated seed failure, with or without pollen coupling, respectively) (Figure 3). However, all simulated time series with among-year variation in resource gain reproduced time series with CV_p_, S, and AC1 similar to observed dynamics (Figure 3).

For white oaks, any mechanism of environmental correlation (i.e. all models with correlated seed production and/or variation in resource gain among years) was sufficient to lead to time series similar to most metrics of observed time series (Figure 3). In other words, the only models that consistently failed were the models with no correlated seed failure and no among-year variation in resource gain.

## Discussion

Correlated seed failure is sufficient to synchronize seed production in general resource budget models, and is likely to be an important factor synchronizing seed production by two species of oaks. Previous theoretical work concluded that environmental noise alone could not drive large-scale spatial synchronization of trees reproduction and pollen coupling is also required (Satake & Iwasa, 2002a,b; Iwasa & Satake, 2004). However, spatially correlated environmental variation is believed to be responsible for the large-scale spatial synchrony in seed production in masting plants (Koenig & Knops, 2000; Satake & Iwasa, 2002b; Koenig & Knops, 2013; Fernandez-Martinez *et al.*, 2017). Our results partly resolve this conflict by including environmental variation in a different way; if correlated environmental noise *per se* is replaced with reproduction failure caused by environmental veto (e.g. drought or late spring frost) then large-scale synchronization might apply without density-dependent pollen limitation. To the extent that an environmental veto synchronizes seed production, the scale of induced synchronization will depend on the scale of the catastrophic event, e.g. late spring-frost has been reported to kill flowers and inhibit reproduction in entire forest communities (Chang-Yang *et al.*, 2016). This would be an interesting hypothesis for further empirical study.

Our results provide mixed support for pollen coupling as a driver of mast-seeding in oaks. Inclusion of density-dependent pollen limitation clearly improved the fit of models to patterns of seed production. However, in simulations, pollen coupling was neither necessary nor sufficient by itself to reproduce patterns of seed production in either species. In general, pollen coupling has been considered unlikely in wind-pollinated species because wind-pollinated plants are able to produce large number of male flowers even when resources are low (Culley *et al.*, 2002; Friedman & Barrett, 2009). Nonetheless, oaks are pollen-limited even in years when flower production is relatively high (Pearse *et al.*, 2015). Thus, environmentally-driven pollination failure in oaks, operating through e.g. phenology synchrony (Koenig *et al.*, 2015; Pesendorfer *et al.*, 2016, Bogdziewicz *et al.*, 2017) could function as a mechanism of correlated seed failure. Our analysis suggests density-dependent pollen limitation may occur in both species, but that pollen coupling is not the primary synchronizing mechanism in oaks. This conclusion is consistent with general theoretical models (Satake & Iwasa 2000; Crone *et al.*, 2005), which indicate that pollen coupling must be sufficiently high, not merely statistically detectable, to synchronize reproduction.

Patterns of seed production were generally similar between the two oak species, in spite of differences in seed development times. In addition, red oak seed production was better predicted by seed resources in the year of seed filling than of pollination, and these non-lagged models were somewhat better at reproducing observed dynamics. Therefore, it seems that the final decision of how much resources will be used to fill acorns happens in the same year in both species. Past studies have shown that the proportion of pollinated flowers is determined the year before seed fall in oaks, but plants still abort significant proportion of acorns in the year of acorn filling, even in mast years (Sork *et al.*, 1993; Espelta *et al.*, 2008; Perez-Ramos *et al.*, 2010, Pearse *et al.*, 2015). The amount of resources invested in acorns may be determined during acorn abscission which happens in the year of acorn maturation, for both red and white oaks (Sork *et al.*, 1993; Owens, 1995; Johnson, 2009). Although our results suggest that red oaks determine seed crop size in the year of seed maturation, the process of resource allocation to reproduction in masting trees is complex (Pearse *et al.*, 2016; Allen *et al.*, 2017), and worthy of further explicit investigation. In nut bearing trees such as *Querucs* anthesis uses carbon remobilized from previous growing seasons (Hoch *et al.*, 2013), but seed maturation uses the current-year carbon (Hoch *et al.*, 2013; Ichie *et al.*, 2013). However, N and P used to build seed tissues probably comes from storage (Alla *et al.*, 2012; Sala *et al.*, 2012; Allen *et al.*, 2017). So, one scenario is that plants first bet how many resources can be spent (anthesis), which is than verified at pollination, and finally by the current-year conditions (Owens, 1995; Fernandez-Martinez *et al.*, 2015; Allen *et al.*, 2017).

To conclude, we showed that correlated seed failure is a powerful synchronizing driver of reproduction in masting plants. Unlike pollen coupling, an environmental veto can synchronize reproduction across species. Unlike pollinator coupling (Tachiki *et al.*, 2010) it could operate in wind-pollinated plants, which comprise significant part of masting species (Herrera *et al.*, 1998; Kelly & Sork, 2002). Reproduction-inhibiting weather events are prevalent in ecosystems (Inouye, 2000), making these mechanisms likely to operate in many masting systems. A final caveat to this study is that, as in any observational study, we were able to rule out possible mechanisms (e.g., pollen coupling in the absence of environmental drivers) than attempt to definitively demonstrate cause and effect. We hope that future studies will investigate the consequences of correlated seed failure more explicitly, using experimental manipulations and/or more detailed studies of physiological mechanisms. Nonetheless, our study resolves a conflict between empirical and theoretical perspectives on the importance of weather events for synchronizing mast seeding (Satake & Iwasa 2002a,b; Koenig & Knops 2013). It also points out the potential importance of changes in extreme events, such as frost or drought, in determining long-term population dynamics.

## Acknowledgments

This work was supported in part by NSF DEB 10-20889 to EEC. MB was supported by the Polish Foundation for Science scholarship ‘Start’, by the (Polish) National Science Foundation grant Preludium no. 2015/17/N/NZ8/01565. MAS recognizes the support of the U. S. National Science Foundation (DEB-1556707).

